# REPEATED INJURY PROMOTES TRACHEOBRONCHIAL TISSUE STEM CELL ATTRITION

**DOI:** 10.1101/2021.01.08.425956

**Authors:** Moumita Ghosh, Cynthia L. Hill, Alfahdah Alsudayri, Scott W. Lallier, Don Hayes, Saranga Wijeratne, John E. Mahoney, Susan D. Reynolds

**Affiliations:** Department of Medicine, University of Colorado-Denver, Denver, CO, USA, 80206; Center for Perinatal Research, Nationwide Children’s Hospital, Columbus, OH, USA, 43215; Department of Pediatrics, The Ohio State University College of Medicine, Columbus, OH; Cystic Fibrosis Foundation Therapeutics, Lexington MA 02421, Cystic Fibrosis Foundation, Bethesda, MD, USA, 20814, USA, 43210

**Keywords:** airway epithelial stem cell, basal cell

## Abstract

Chronic lung disease has been attributed to stem cell aging and/or exhaustion. To address this issue, we investigated the lifespan of tracheobronchial tissue stem cells (TSC) over time and in response to repeated injury. Chromatin and nucleotide labeling studies in mice indicated that: 1) injury activated a subset of the TSC pool and that this process conserved TSC over time; and 2) activated TSC were predisposed to further proliferation and this activated state lead to terminal differentiation. Analysis of human TSC and clonal isolates indicated that repeated TSC proliferation led to telomere shortening and analysis of TSC from Dyskeratosis Congenita donors indicated that mutations in telomere biology genes accelerated TSC depletion. RNAseq and functional studies indicated that human TSC terminated as a secretory committed cell. These data support a model in which a repeated epithelial injury depletes the TSC pool and initiates the abnormal repair associated with chronic lung disease.

## INTRODUCTION

A tissue specific stem cell (TSC) is defined as a cell that self-renews and generates each of the differentiated cell types found in the stem cell’s home tissue. Clonal analysis in human (Engelhardt, Schlossberg et al. 1995) and lineage-tracing in mice (Hong, Reynolds et al. 2003, Hong, Reynolds et al. 2004, Rock, Onaitis et al. 2009, Ghosh, Brechbuhl et al. 2011) indicated that the tracheobronchial TSC was a basal cell subtype. This TSC generated basal, ciliated, and secretory epithelial cells as well as a variety of rare epithelial cell types.

Conducting airway epithelial remodeling is a hallmark of chronic lung diseases including cystic fibrosis, asthma, chronic obstructive pulmonary disease (COPD), and idiopathic pulmonary fibrosis (IPF) (Jeffery 2001). Focal lesions are associated with repeated injuries and an aberrant epithelial repair process that alters the frequency of differentiated epithelial cells. Since these differentiated cells are derived from the TSC, epithelial remodeling has been attributed to TSC aging and/or exhaustion (Armanios and Blackburn 2012). While abnormal TSC function (i.e. a change and self-renewal and/or multi-lineage differentiation) and/or TSC depletion may underlie epithelial pathology, specific cellular and molecular mechanisms have not been established.

We previously reported that mouse TSC generated a specific clone type, the rim clone, in vitro (Ghosh, Helm et al. 2011). Serially passaged rim clones terminated when they generated a non-rim clone. These non-rim clones were composed of non-mitotic basal cells. Thus, the last mouse TSC-derived progenitor cell was the unipotential basal (UPB) cell.

We used flow cytometry and clone morphology to show that naphthalene (NA)-mediated airway epithelial injury in mice activated approximately half the stem cell pool (Ghosh, Helm et al. 2011). TSC activation resulted in a transient increase in TSC number and TSC frequency was normalized via a terminal differentiation process. To determine if mitotic frequency was associated with terminal differentiation, we used the Histone 2B:Green Fluorescent Protein (H2B:GFP) chromatin labeling method (Tumbar, Guasch et al. 2004) to purify TSC that proliferated at different rates (Ghosh, Smith et al. 2013). TSC that proliferated infrequently in response to NA injury generated significantly more rim clones than TSC that proliferated frequently. Finally, serial passage analyses demonstrated that repeated cell division drove the terminal differentiation process. These studies challenged the idea that the TSC maintains its function throughout an individual’s life and suggested a proliferation-based aging mechanism.

Aging, particularly at the cellular level, has been associated with telomere shortening (Armanios and Blackburn 2012). Telomeres are 6 base pair repeats that form the ends of chromosomes. Telomere repeats are generated by the reverse transcriptase telomerase (TERT). Mouse telomeres are ∼200 kilobases (kb) in length; whereas human telomeres are ∼25-50 kb, (Greider 1996)). Consequently, telomere length studies are frequently conducted in human or in mice which harbor TERT mutations.

Telomeres prevent loss of genetic information during DNA replication (Armanios and Blackburn 2012). In parallel, the shelterin protein complex interacts with telomere repeats and prevents recognition of the single-stranded chromosome end as a double-strand break. Most mammalian somatic cells express little or no TERT, consequently telomeres shorten with each cell division. In vivo and in vitro studies demonstrated that telomere length was inversely correlated with age and that critically short telomeres led to cell cycle arrest. Whether the arrested cell becomes terminally differentiated, senescent, apoptotic, or transformed is dependent on additional factors.

Dyskeratosis Congenita (DC) is caused by germline mutations in telomere biology genes including TERT and genes that encode the various shelterin complex proteins (Higgs, Crow et al. 2019). DC patients present with nail dystrophy, changes in skin pigmentation, and oral leukoplakia. The most severe forms of DC lead to bone marrow failure. Analysis of telomere length in leukocytes associated telomere shortening with hematopoietic stem cell attrition and premature aging. The dominant effect of telomere biology gene mutations allowed us to determine if loss of function altered TSC number or life span.

Based on our previous report that NA injury activated half the TSC pool, this project evaluated the behavior of the activated and quiescent TSC subsets as a function of time and in response to a second injury. To investigate the mechanism underlying TSC attrition, we stimulated TSC proliferation through serial passage of human TSC from normal and DC patients. TSC number, lifespan, differentiation potential, and telomere length were quantified.

## RESULTS

### TSC continue to proliferate after the epithelium is repaired

This study determined the mitotic frequency of TSC following a single NA exposure. K5/H2B:GFP mice and mono-transgenic controls were fed Dox chow for 3 days, treated with NA on day 0, and switched to standard chow on day 3 (Fig 1A). Mice were recovered to day 6, 40, or 80. Flow cytometry was used to evaluate GFP fluorescence intensity (Fig 1B). The frequency of GFP-Bright, -Dim, and - Negative cells within the CD49f-Bright/Sca1+ basal cell population was quantified across 3 experiments (Fig 1C-E).

**Fig 1.**
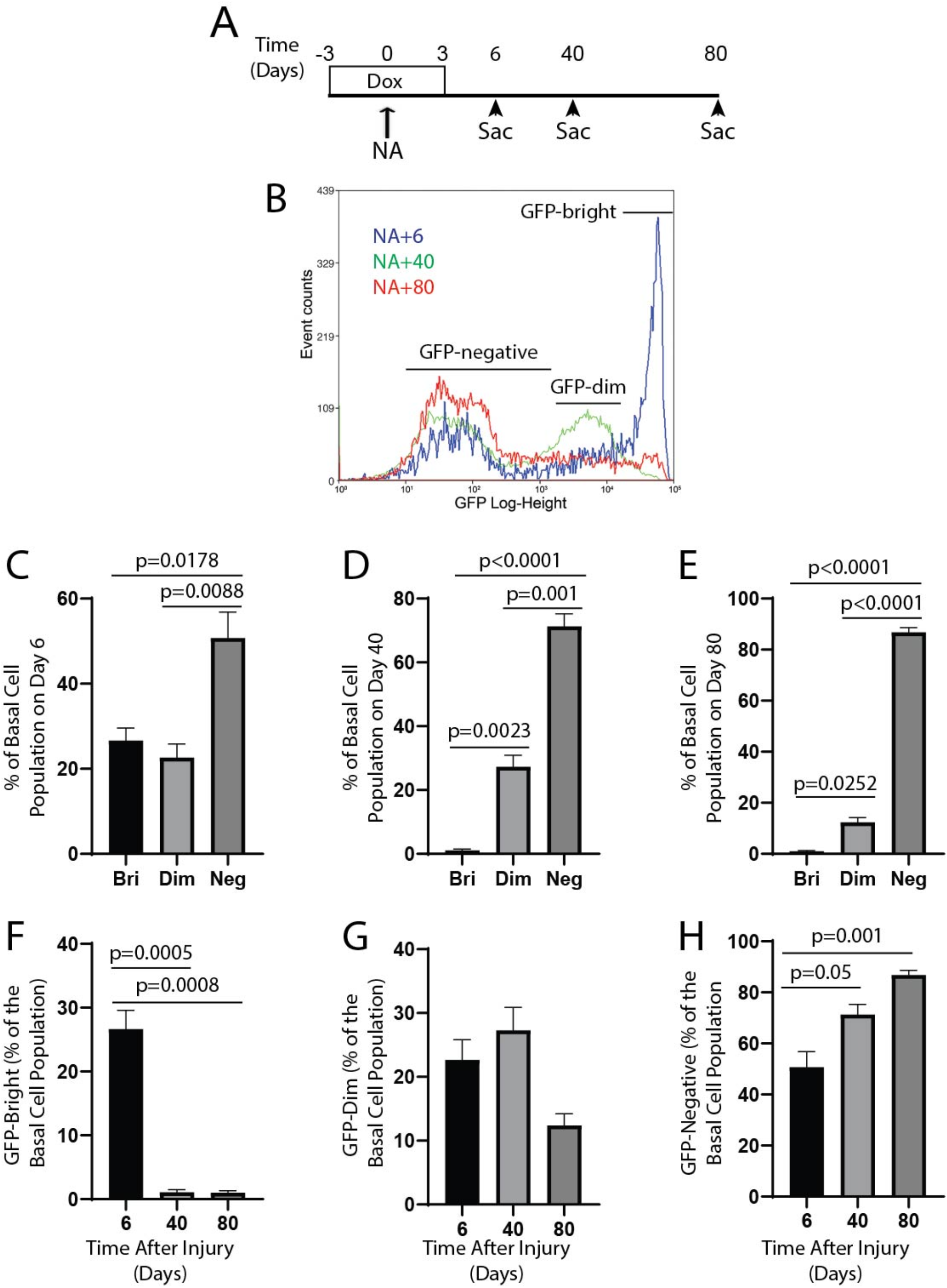
Analysis of TSC cell cycle frequency after a single NA injury. **A**. Experimental design. K5/H2B:GFP mice and mono-transgenic controls were treated with Dox chow from day −3 through day +3 and then switched to standard chow. NA treatment occurred on day 0. Animals were euthanized (sac) on days 6, 40, and 80. **B**. Representative flow cytometric analysis of green fluorescent protein (GFP) fluorescence intensity. **C-E**. Frequency of GFP-Bright (Bri), GFP-Dim, and GFP-Negative (Neg) cells on days 6 (C), 40, (D), and 80 (E). **D-F**. Frequency of GFP-Bri (F), GFP-Dim (G), and GFP-Neg (H) a function of time after injury. Mean ± SEM (n=3).

The frequency of GFP-bright cells decreased significantly between days 6 and 40 but did not change between days 40 and 80 (Fig 1F). These data indicated that the GFP-Bright population stabilized between days 40 and 80. The frequency of GFP-Dim cells did not vary as a function of time (Fig 1G). However, the frequency of GFP-Negative basal cells was significantly increased on days 40 and 80 (Fig 1H). These data indicated that a single NA injury generated a GFP-Bright population and that most of these cells proliferated by 80.

A previous study reported that a 2-fold decrease in GFP mean fluorescence intensity (MFI) occurred with each cell division (Blanpain, Lowry et al. 2004). The MFI was 6×10^4^ for GFP-Bright cells, 5×10^3^GFP-Dim cells, and 4×10^3^ for GFP-Negative cells (Fig 1B). These data indicate that many GFP-Bright cells divided ∼3 times between days 6 and 40 and another 3 times between days 40 and 80. Knowing that the tracheal epithelium is repaired within 13 days of NA injury (Cole, Smith et al. 2010) and that most GFP-Bright cells are TSC (Ghosh, Smith et al. 2013) these data indicated that TSC continued to proliferate after the epithelium was repaired.

### A second injury accelerates TSC proliferation

This study focused on TSC that proliferated in response to the first NA injury and retained the chromatin label to day 40. To determine if a second injury altered their mitotic activity, K5/H2B:GFP mice and controls were fed Dox chow and treated with NA on day 0. The mice were switched to standard chow on day 3 and recovered to day 40 (Fig 2A, top panel). A second group of mice was retreated with NA on day 40 and recovered to day 80 (Fig 2A, bottom panel). Flow cytometry was used to evaluate GFP fluorescence 40 days after the first or second injury.

**Fig 2.**
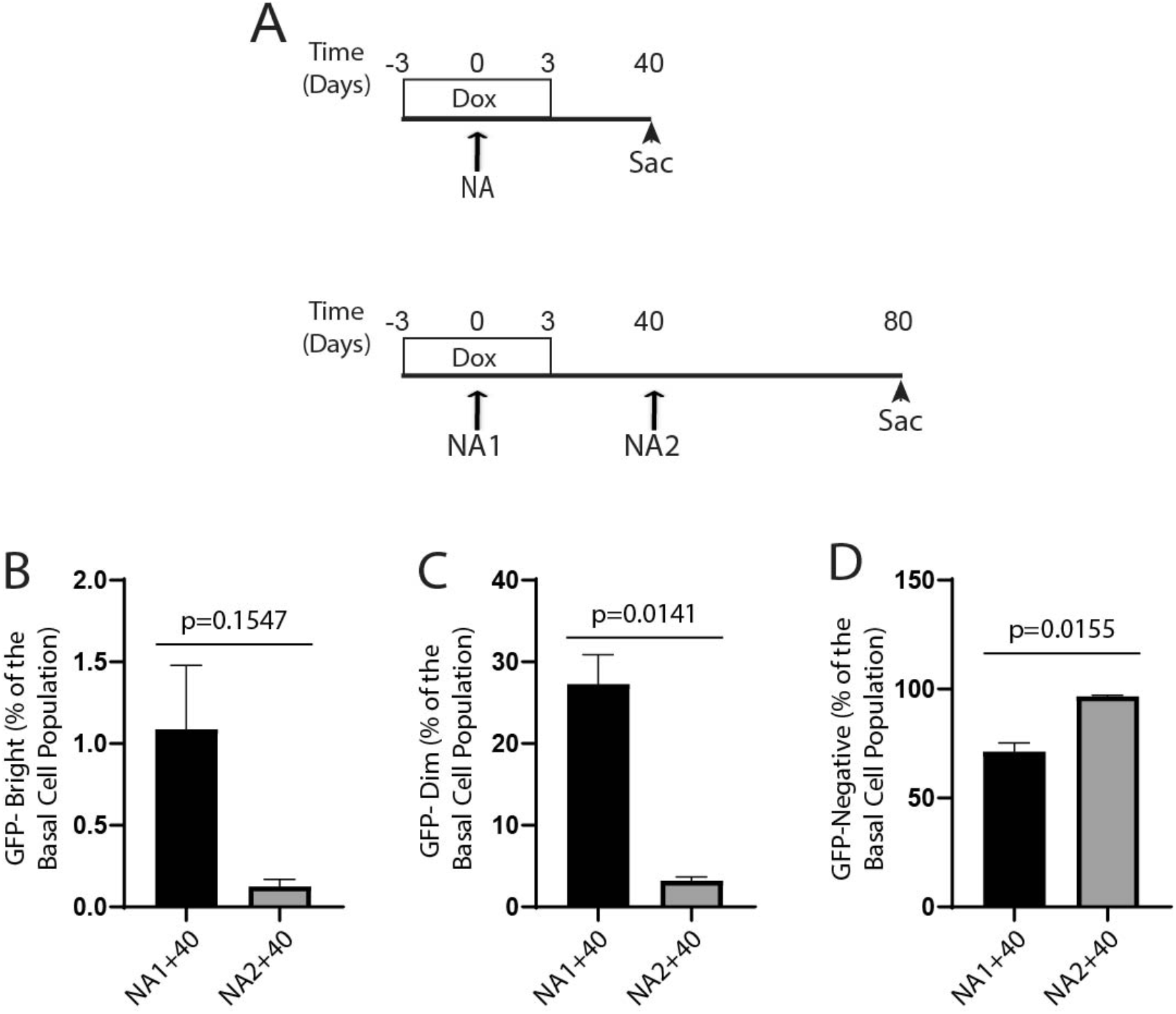
Analysis of TSC cell cycle frequency after a second NA injury. **A**. Experimental design. Top Panel. K5/H2B:GFP mice and mono-transgenic controls were treated with Dox chow from day −3 through day +3 and then switched to standard chow. NA treatment occurred on day 0. Animals were euthanized (sac) on day 40. Bottom Panel. Mice were treated as indicated in the Top Panel and retreated with NA on day 40 (NA2). Mice were euthanized on day 80. Flow cytometry was used to evaluate green fluorescent protein (GFP) fluorescence intensity 40 days after NA1 or NA2. **B**. Frequency of GFP-Bright cells 40 days after NA1 or NA2. **C**. Frequency of GFP-Dim cells 40 days after NA1 or NA2. **D**. Frequency of GFP-Negative cells 40 days after NA1 or NA2. Mean ± SEM (n=3).

The frequency of GFP-Bright cells did not change significantly in response to the second injury (Fig 2B). However, the MFI for GFP-Bright cells decreased (SFig 1). The frequency of GFP-Dim cells (Fig 2C) was significantly decreased after the second injury and the frequency of GFP-Negative cells increased (Fig 2D). This analysis indicated that GFP-Bright cells proliferated in response to the second injury and the second injury accelerated the proliferation of TSC within the GFP-Dim population. These data suggested that the initial injury/repair process generated a pool of “activated” TSC which were predisposed to proliferate in response to the second injury.

### Terminal differentiation depletes the TSC pool

Our previous studies demonstrated that TSC have a finite life-span and that proliferation in vivo or in vitro promoted terminal differentiation to a UPB cell (Ghosh, Helm et al. 2011, Ghosh, Smith et al. 2013). In the present study we used a cloning strategy to investigate differentiation of TSC which had proliferated in response to NA injury. K5/H2B:GFP mice were treated as indicated above (Fig 1A). On day 40, TSC were recovered, cloned, and serially passaged. This analysis demonstrated that TSC recovered from NA-injured mice survived at least 3 passages (SFig 2). An analysis of rim (TSC-derived clones) and non-rim (UPB-derived clones) showed that clones 1 and 2 generated significantly more TSC clones than UPB cell clones. In contrast, clones 3 and 4 generated significantly fewer TSC clones than UPB cell clones. Clone 4 terminated at passage (P) 3.

To further evaluate the behavior of TSC, limiting dilution was used to generate clonal sublines from the three long-lived TSC clones (Clones 1-3). The clone-1 sublines generated only TSC clones at P4 (Fig 3A). Sub-cloning generated 40 clones at P5 (Fig 3B). Two of the P5 sub-clones (5%) generated only TSC clones; whereas, the remaining 38 clones (95%) generated a mixture of TSC and terminally differentiated UPB clones. Each of the mixed sub-clones generated more TSC clones than UPB clones.

**Fig 3.**
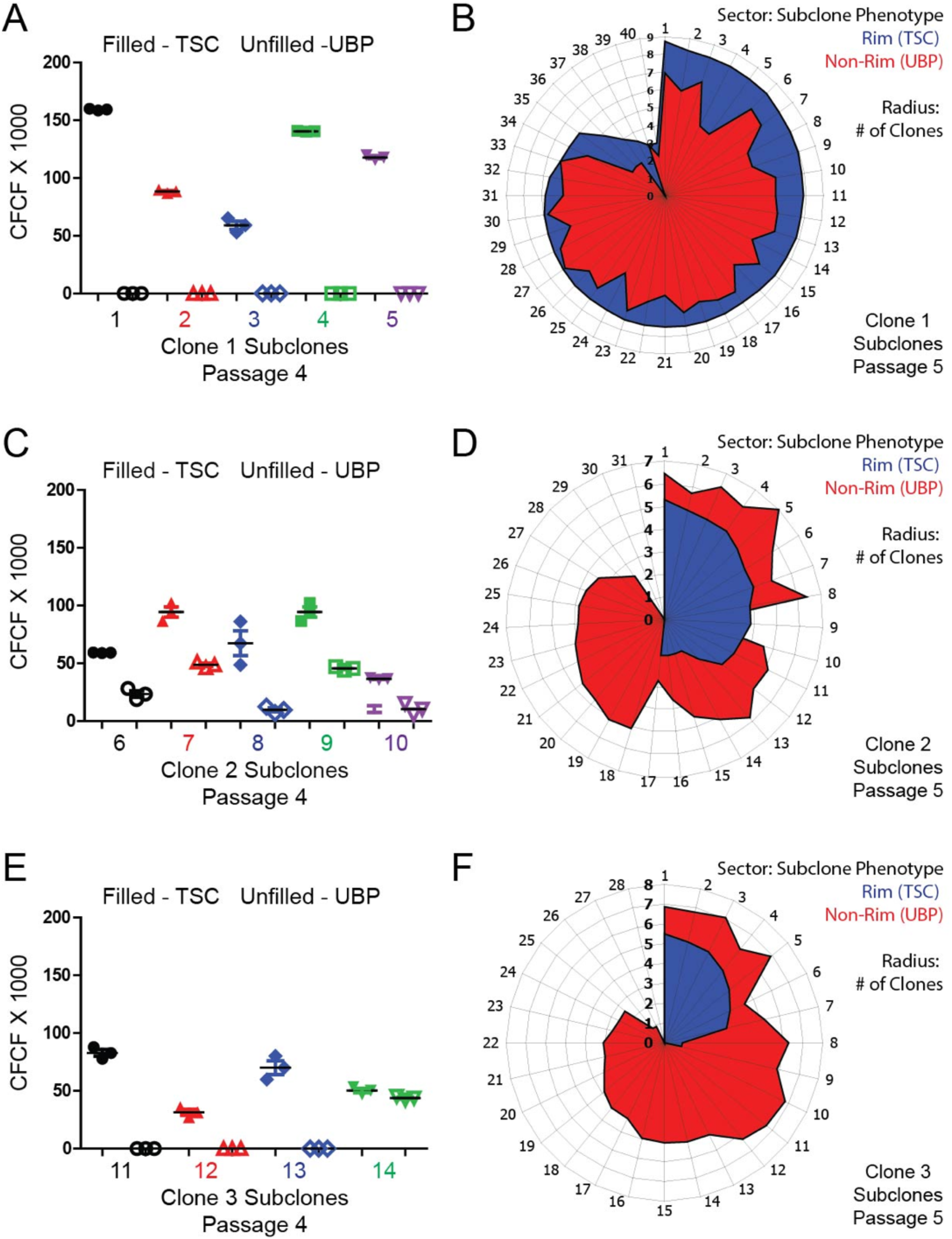
Lifespan and fate analysis of TSC clones. K5/H2B:GFP mice and mono-transgenic controls were treated with dox chow from day −3 through day +3 and then switched to standard chow. NA treatment occurred on day 0. On day 40, TSC were recovered, cloned, and serially passaged. Long-lived clones, numbers 1-3 (see SFig 2 for details) were subcloned at passages 2, 3, and 4. **A-B**. Clone 1, subclones 1-5. **C-D**. Clone 2, subclones 6-10. **E-F**. Clone 3, subclones 11-14. The fifth clone did not survive. **A, C, E**. TSC (filled symbols) and UPB (open symbols) clone forming cell frequency (CFCF) analysis for subclones at passage 4. **B, D, F**. The number of TSC and UPB colonies per TSC clone at passage 5. Sectors represent subclones. Diameter represents the number of TSC (blue) or UPB (red) clones.

Clone-2 sublines generated both TSC and UPB clones at P4 (Fig 3C). Sub-cloning of 31 TSC identified 29 clones (94%) that survived to P5 (Fig 3D). Among the surviving TSC clones, 2 (6.9%) generated only TSC, 14 (48%) generated fewer TSC clones than UPB clones, and 12 (41%) generated only UPB clones.

Three Clone-3 sublines generated only TSC clones at P4, while the fourth subline generated both TSC and UPB clones (Fig 3E). Sub-cloning of 28 TSC identified 27 clones (96%) that survived to P5 (Fig 3F). Seven of these clones (26%) generated both TSC and UPB clones; whereas, 18 clones (67%) generated only UPB.

This analysis demonstrated that only 4.3% of TSC clones self-renewed at P5 and that the vast majority of TSC had entered the terminal differentiation pathway. Collectively, this analysis of cloned TSC indicated that repeated proliferation and terminal differentiation depleted the activated TSC pool.

### Comparable TSC proliferation within 6 days of the first and second NA injuries

Injury-induced TSC depletion caused us to compare the mitotic frequency of TSC after the first and second NA exposures. Group 1 consisted of K5/H2B:GFP mice and controls which were fed Dox chow for 3 days, treated with NA on day 0, switched to standard chow on day 3, and recovered to day 6 (Fig 4A). Group 2 consisted of K5/H2B:GFP mice and controls which were treated with NA on day 0, switched to Dox chow on day 37, retreated with NA on day 40, switched to standard chow on day 43, and recovered to day 46 (Fig 4B). Analysis of GFP fluorescence demonstrated that Group 2 contained GFP-bright cells and that the pattern of GFP fluorescence was indistinguishable from that observed in Group 1 (SFig 3A). Flow analysis of epithelial cell phenotype in Groups 1 and 2 did not detect differences in the frequency of epithelial, basal, GFP-Dim, or GFP-Bright cells (Fig 4D). These data indicate that the early TSC response to the first and second injuries were similar.

**Fig 4.**
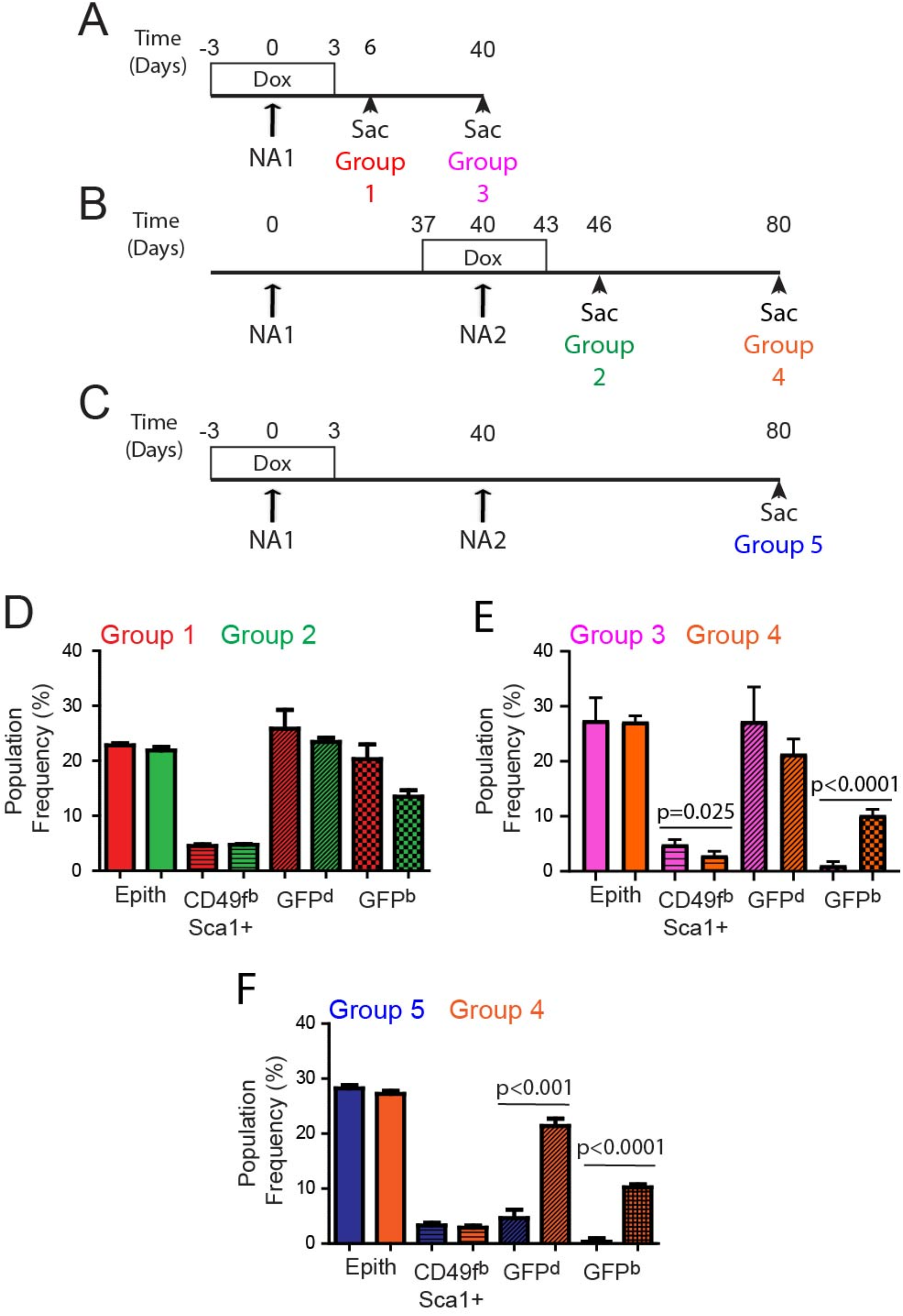
Comparison of the TSC response to a single or repeated NA injury. **A-C**. Experimental design. All studies used K5/H2B:GFP mice and mono-transgenic controls. **A**. Mice were treated with Dox chow from day −3 through day +3 and then switched to standard chow. NA treatment occurred on day 0. Animals were euthanized (sac) on days 6 (Group 1) or 40 (Group 3). **B**. Mice were treated with NA on day 0, fed Dox chow from days 37-43, and retreated with NA on day 40 (NA2). Mice were euthanized on days 46 (Group 2) and 80 (Group 4). **C**. Mice were treated with Dox and NA as indicated in panel A and retreated with NA on day 40 (NA2), and recovered to day 80 (Group 5). **D**. Frequency of epithelial (Epith), basal cells (CD49f^b^/Sca1^+^), GFP-Dim (GFP^d^), and GFP-Bright (GFP^b^) cells in Groups 1 and 2. The frequency of each subset within the CD45-/CD31-/TER119-/DAPI-population was determined. Mean ± SEM (n=3). **E**. Frequency of Epith, CD49f^b^/Sca1^+^, GFP^d^, and GFP^b^ cells in Groups 3 and 4. Mean ± SEM (n=3). **F**. Frequency of Epith, CD49f^b^/Sca1^+^, GFP^d^, and GFP^b^ cells in Groups 4 and 5. Mean ± SEM (n=3).

### Divergent TSC proliferation 40 days after the first and second NA injuries

To compare TSC mitotic frequency at later time points, we evaluated GFP fluorescence and cell phenotype 40 days after the first or second NA injury. Group 3 mice were treated with Dox and NA as indicated for Group 1 and were recovered to day 40 (Fig 4A). Group 4 mice were treated with Dox and NA as indicated for Group 2 and were recovered to day 80 (Fig 4B). Analysis of GFP fluorescence and epithelial phenotype demonstrated that Group 3 had significantly more CD49f-Bright/Sca1+ cells than Group 4 and that Group 3 had significantly fewer GFP-Bright cells than Group 4. These data indicated that Group 3 cells proliferated more frequently than Group 4 cells. These data suggested that two mitotically-distinct populations of TSC were involved in epithelial repair.

To further evaluate this finding, we compared GFP fluorescence and cell phenotype in Group 4 mice with K5/H2B:GFP mice and controls which were fed Dox chow for 3 days, treated with NA on day 0, switched to standard chow on day 3, retreated with NA on day 40, and recovered to day 80 (Group 5, Fig 4C). Analysis of GFP fluorescence showed distinct profiles for Groups 4 and 5 (SFig 3B). Quantification demonstrated that the frequency of epithelial and basal cells did not differ for Groups 4 and 5 (Fig 4F). However, the frequency of GFP-Dim and GFP-Bright cells was significantly decreased in Group 5 relative to Group 4. These data further supported the idea that cells labeled during the first injury continued to proliferate after the second injury and raised the possibility that the second injury activated a distinct cohort of TSC.

### The second injury activates a new cohort of TSC

To determine if the same set of TSC responded to the first and second injuries, we used the nucleotide label-retention technique. Mice were treated with NA on day 0 and mitotic cells were labeled with BrdU (Fig 5A, upper panel). The frequency of BrdU label-retaining cells was assayed on days 40 and 80. A group of mice was re-injured and mitotic cells were labeled with EdU. The frequency of EdU+, BrdU+, and BrdU+/EdU+ label-retaining cells was assayed on day 80 (Fig 5A, lower panel).

**Fig 5.**
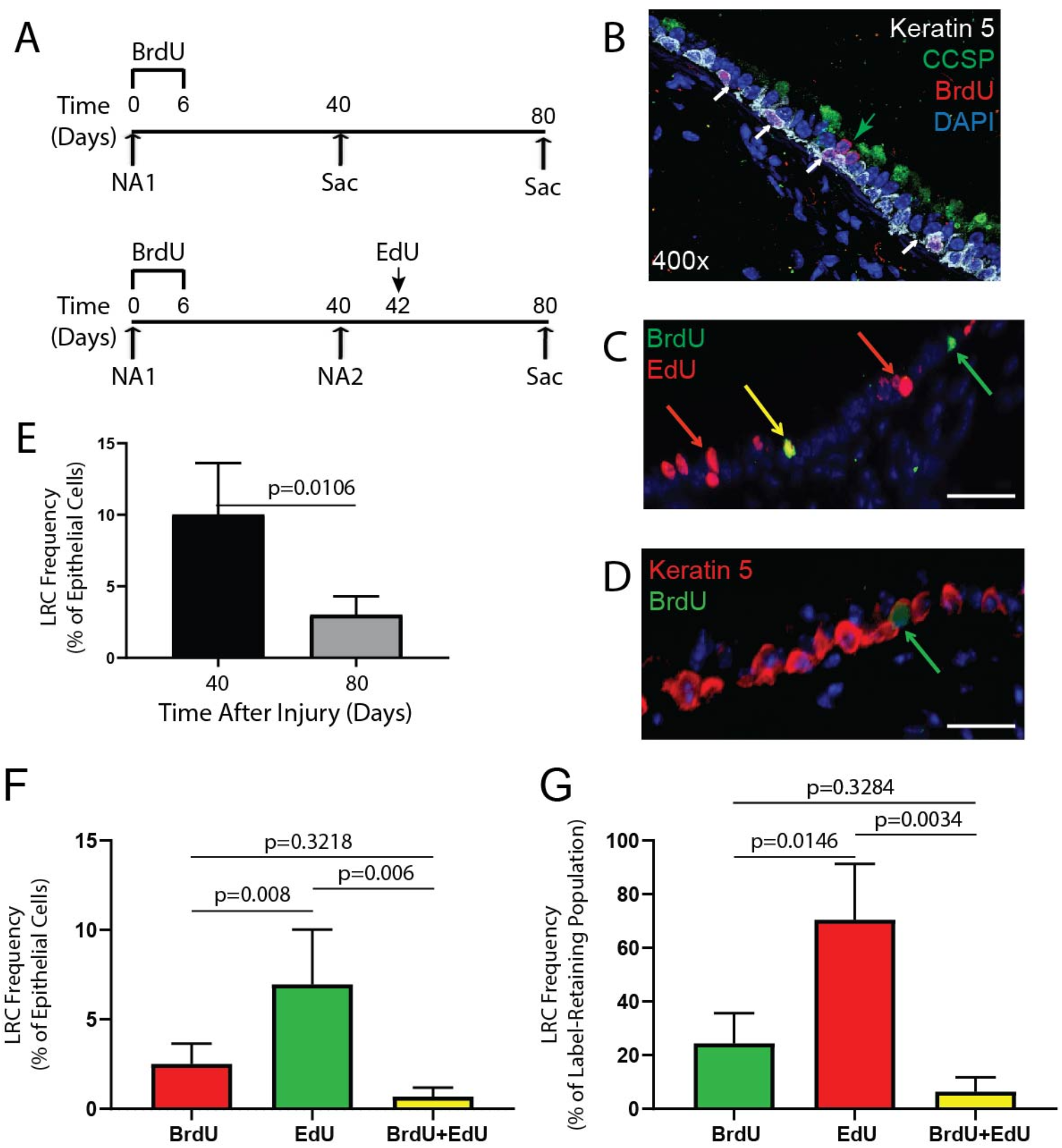
Nucleotide label retention analysis of cells that proliferate after the first and second NA injuries. **A**. Experimental design. Top Panel. FVB/n mice were treated with NA on day 0 and Bromo-deoxyuridine (BrdU) on days 0-6. Tissue was recovered on day 40 (sac). Bottom Panel. Mice were treated as indicated in the Top Panel (NA1) and retreated with NA on day 40 (NA2). Mice were treated with 5-ethynly-2’-deoxyuridine (EdU) on day 42. Tissue was recovered on day 80. **B**. Histological analysis of Keratin 5 (white), CCSP (green), BrdU (red), and DAPI (blue) 40 days after NA injury. White arrows indicate Keratin 5/BrdU+ cells. **C**. Histological analysis of BrdU (green) and EdU (red) on day 80. Green arrow indicates a BrdU+ cell. Red arrows indicate EdU positive cells. Yellow arrow indicates a BrdU/EdU positive cell. Scale bar, 50 microns. **D**. Histological analysis of Keratin 5 (red) and BrdU (green) on day 80. Green arrow indicates a Keratin 5/BrdU double-positive cell. Scale bar, 50 microns. **E**. Frequency of BrdU label retaining cells (LRC) within the epithelial cell population on recovery days 40 and 80. **F-G**. Quantification of BrdU+, EdU+, and BrdU+/EdU+ LRC on day 80. **F**. Frequency of BrdU+, EdU+, and BrdU+/EdU+ LRC as a percentage of all epithelial cells. **G**. Frequency of BrdU+, EdU+, and BrdU+/EdU+ LRC as a percentage of the label-retaining cell population. **E-G**. Mean ± SEM (n=3-5).

Histological analysis (Fig 5B-D) and quantification demonstrated that the frequency of BrdU+ cells decreased significantly between days 40 and 80 (Fig 5E). These data are consistent with the chromatin labeling studies (Fig 2) and indicate that many BrdU-tagged cells proliferated in response to the second injury. On Day 80, the frequency of EdU+ cells was significantly greater that the frequency of BrdU+ and BrdU+/EdU+ cells and the frequency of BrdU+ and BrdU+/EdU+ cells was similar (Fig 5F). Further, the frequency of EdU+ cells within the label-retaining cell population was significantly greater than that of the BrdU+ or BrdU+/EdU+ populations (Fig 5G). These data supported the idea that the second injury activated a new cohort of TSC.

### Telomere shortening in serially passaged human TSC

TSC depletion in mice resembled the loss of clone forming cells in serially-passaged human TSC ((Hayes, Kopp et al. 2018), Fig 6A) and raised the possibility that short-lived TSC had proliferated repeatedly prior to isolation. Since proliferation of somatic cells and TSC attrition has been associated with telomere shortening (Goodell, Rosenzweig et al. 1997), we used PCR to determine telomere length in serially passaged human TSC. This study demonstrated that telomeres shortened as a function of passage (Fig 6B). To determine if serial passage was selecting TSC with relatively long telomeres, human TSC were cloned at P3 and telomere length was determined for the parental populations and clonal isolates at P4-P11. Telomere length was significantly greater in clones than in populations at P4 to P6 (Fig 6C). These data indicate that short-lived TSC enter culture with relatively short telomeres and that an additional decrease in telomere length contributes to TSC attrition over P1 through P5.

**Fig 6.**
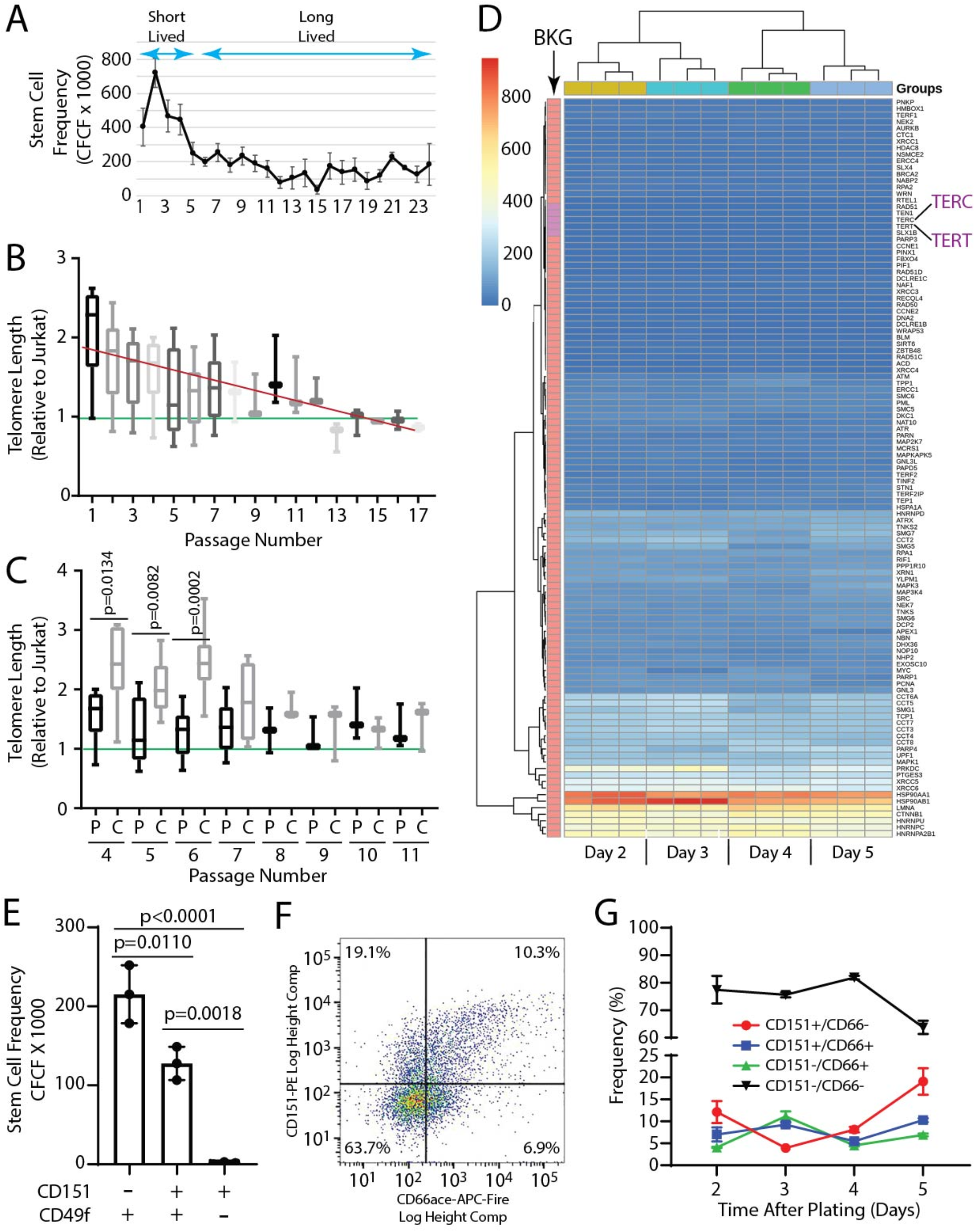
Telomere length and secretory differentiation in human TSC. **A**. The clone forming cell frequency (CFCF) assay was used to determine TSC number across 24 passages. Mean ± SD (n=4 donors). **B**. Telomere length in TSC from normal donors. Green line indicates telomere length that is equal to that of the short telomere control, Jurkat. Red line indicates the linear regression analysis of telomere length and passage. Min-Max graphs with line at mean, box indicates 10-90 percentile (n=5-9 for passages 1-7; n=3 for passages 8-17). **C**. Telomere length in populations of TSC and clonal TSC isolates. Green line indicates telomere length that is equal to that of Jurkat. Min-Max graphs with line at mean, box indicates 10-90 percentile (n=9 for populations; n=7 for clones). **D**. Heatmap illustrating RNAseq analysis of passage 2 TSC on post-plating days 2, 3, 4, and 5 (n=3). Expression, in Fragments Per Million Reads (FPM), for telomere biology genes is shown. Background (BKG): FPM > 0.5 = above background (salmon), FPM<0.5 below background (plum including TERT and TERC). **E**. Clone forming cell frequency in TSC subsets defined by expression of CD49f and CD151. Mean ± SEM (n=3). **F**. Representative flow cytometry analysis of CD151 and CD66ace expression in long-lived TSC. Live/CD49f+ cells were analyzed. **G**. Frequency of CD151 and CD66ace within the live/CD49f+ population. Changes as a function of time after plating are shown. Day 2, ∼10% confluent; Day 3, ∼40% confluent, Day 4, ∼80% confluent; Day 5, ∼99% confluent. Mean ± SEM (n=3).

The telomere length study also found that telomere length was stable, but short, at later passages (Fig 6C). To determine if long-lived TSC expressed telomere maintenance genes TERT and TERC, we used RNAseq to evaluate expression of these genes as a function of time after plating. This study did not detect TERT or TERC mRNA at any time point (Fig 6D, SFig 4) and was in agreement with other studies did not detect TERT gene expression or function (Mou, Vinarsky et al. 2016, Reynolds, Rios et al. 2016). However, TERT activity has been reported in TSC (Yim, Slebos et al. 2007, Suprynowicz, Upadhyay et al. 2012) and it was possible that these population studies may have overlooked TERT and TERC expression in a rare subpopulation. To address this issue, we queried a single cell RNAseq data sets for expression of telomere biology genes (bioRxiv 2020.05.01.072876). Analysis of freshly isolated human lung cells as well as TSC that were proliferating in vivo did not detect a TERT- or TERC-expressing subpopulation (SFig 5). Collectively, these data indicate that TSC do not express TERT or TERC.

### Secretory specification stabilizes TSC frequency

An additional characteristic of long-lived human TSC was stabilization of clone forming cell frequency at 5-10% (Fig 6A). Our analysis of mouse TSC clones (Fig 3) indicated that most TSC generated a daughter TSC and a terminally differentiated UPB cell. In contrast, we previously noted that human TSC upregulated secretory genes as they were passaged (Reynolds, Rios et al. 2016). To further assess this observation, we used RNAseq to evaluate expression of the secretary cell type signature (Carraro, Mulay et al. 2020) in P2 TSC as a function of time after plating (SFig 6). This study indicated that expression of most secretory signature genes increased as the cultures approached confluence.

To further evaluate secretory differentiation, we first evaluated CD151, a secretory commitment marker (Ghosh, Ahmad et al. 2013). An analysis of CFCF indicated that CD151+ cells had significantly fewer clone forming potential than CD151-cells (Fig 6E). Next, we evaluated expression of CD151 and the secretory priming marker CD66ace (Carraro, Mulay et al. 2020) and demonstrated that CD151+ cells were a subset of the CD66+ population (Fig 6F). Finally, we quantified the frequency of the primed and committed subtypes. This study indicated that secretory primed cells were generated early in culture while secretory committed cells accumulated as the cultures approached confluence. These data indicated that a subset of TSC daughter cells commitment to the secretory differentiation pathway and that this process limited the frequency of clone forming human TSC.

### TSC number and function are decreased in DC

In contrast with TERT and TERC, our RNAseq study detected expression of shelterin genes including DKC1, NAF1, NOT10, TERF1, TERF2, TPP1, and WRAP53 (Fig 6D) and raised the possibility that long-lived TSC used this complex to protect their relatively short telomeres. To address this question, we evaluated TSC from donors who harbor mutations in TERT or shelterin genes. The study group contained 12 DC patients and 16 parents (STable 1).

Within Family 1, the mother and the proband (DC patient) harbored a mutation in RTEL1. A previous study reported telomere shortening in their peripheral blood leukocytes (Ballew, Yeager et al. 2013). The father exhibited premature aging and telomere shortening but a specific mutation was not identified. The parents and the proband had significantly fewer TSC than control at P1 and the proband had fewer TSC than the parents (Fig 7A). A comparison of 12 DC probands and 16 parents demonstrated that DC probands had fewer TSC than their adult relatives at P1 (Fig 7B).

**Fig 7.**
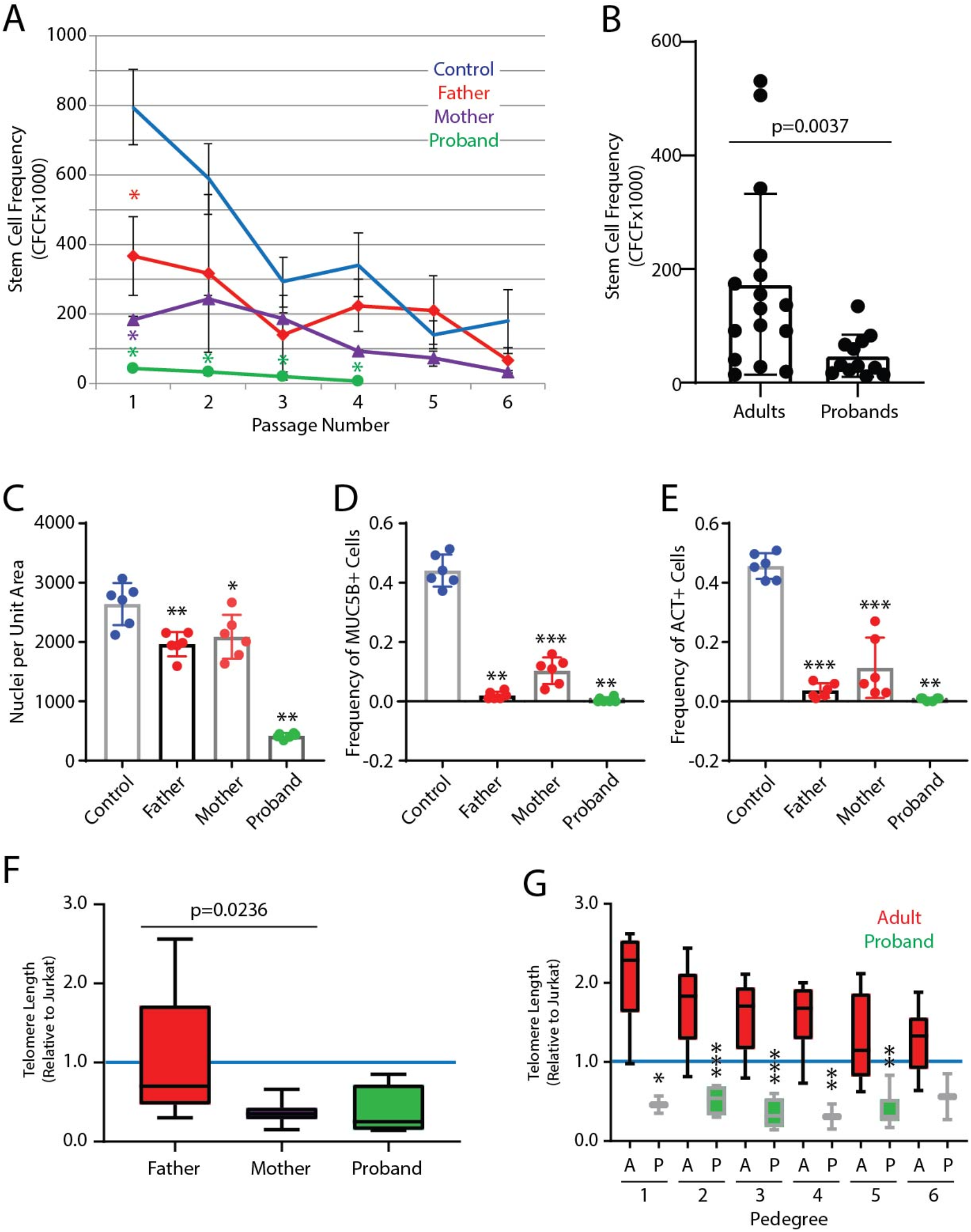
TSC number and function in Dyskeratosis Congenita (DC). **A**. The clone forming cell frequency (CFCF) assay was used to determine TSC frequency in nasal respiratory epithelial cell isolates from normal controls and in Family 1 (see STable 1 for details). Red, purple, and green asterisks indicate significant differences from control (p<0.05). Mean ± SD (n= 3 determinations/donor). **B**. The CFCF assay was used to determine TSC frequency in nasal respiratory epithelial isolates from 12 DC families (see STable 1 for details). **C-E**. TSC differentiation to functional mucus and multiciliated cells was assayed via the air-liquid-interface method. **C**. Nuclear density as assessed by DAPI-positive cell area. **D**. Mucus cell differentiation as assessed by MUC5B-positive cell frequency. **E**. Ciliated cell differentiation as assessed by acetylated tubulin (ACT)-positive cell frequency. Mean ± SD (n=3 replicates/donor). * p<0.05, ** p<0.01, *** p<0001. **F**. Telomere length in TSC from Family 1. Blue line indicates telomere length that is equal to that of the short telomere control, Jurkat. Min-Max graphs with line at mean, box indicates 10-90 percentile (n=3 determinations/donor). **G**. Telomere length in TSC from 12 DC families. Blue line indicates telomere length that is equal to that of Jurkat. Min-Max graphs with line at mean, box indicates 10-90 percentile (n=3 determinations/donor). * p<0.05, ** p<0.01, *** p<0001.

Serial passage of Family 1 TSC demonstrated that TSC were depleted more rapidly in the parents and proband than in control (Fig 7A). TSC from the proband terminated at P4. Similar results were obtained for the other DC Families (data not shown). The ability of TSC from Family 1 to generate functional secretory and ciliated cells was quantified at P2 using air-liquid-interface cultures. Cultures from all donors polarized and indicated that TSC were present. However, cell density and the frequency of goblet cells and ciliated cells was significantly less than control (Fig 7C-E). Similar results were obtained for additional DC families (data not shown). These data indicate that mutations in telomere biology genes, including TERT and shelterin genes, decrease TSC number and function.

### TSC from DC patients have abnormally short telomeres

An analysis of telomere length in TSC from Family 1 demonstrated that the father had significantly longer telomeres than the mother and that telomere length in TSC from the mother and the proband were similar (Fig 7F). An analysis of additional DC families demonstrated that telomeres were significantly longer the parents relative to the proband in five of six comparisons (Fig 7G). These data extend the previously reported telomere shortening in DC-leukocytes to include airway epithelial TSC and supported the idea that shelterin genes are critical for protection of short telomeres in long-lived TSC.

## DISCUSSION

Our previous study indicated that TSC, like other progenitor cell types, obeyed the Hayflick limit (Hayflick 1979). Thus, TSC lifespan was limited to ∼40-50 population doublings. TSC are thought to conserve their mitotic potential by limiting the number of times they divide in response to a stimulus. Based on our finding that injury activated a subset of TSC (Ghosh, Helm et al. 2011), we suggested that partial activation of the TSC pool also conserved TSC. These data further support the idea that selective activation of the TSC pool insures epithelial repair over multiple injuries. However, we suggest that each injury/repair cycle decreases the reparative potential of the epithelium and that the magnitude of this decrease is dependent on the number of TSC that are activated by the injury.

The idea that telomere shortening limits TSC life-span is supported by studies in other systems (Armanios and Blackburn 2012). We associated TSC depletion with repeated proliferation (Figs 1-5) and we now report that these phenomena are paralleled by telomere shortening (Fig 6, 7). Differences in telomere length for TSC populations and clones (Fig 6) indicate that short-lived human TSC have relatively short telomeres. This conclusion is supported by our finding that TSC frequency was significantly decreased in DC patients relative to non-DC controls, long-lived TSC were not detected in DC patients, and TSC from DC patients have short telomeres (Fig 7).

Collectively, our data indicate that telomere shortening occurs as TSC divide and that differences in telomere length reflect the number of times a TSC has undergone cytokinesis. Based on these findings and our analysis of mouse TSC (Figs 1-5), we suspect that freshly isolated human TSC contain subsets that are defined by the number of times they proliferated in vivo. TSC which are lost over passages 1-5 are likely to be cells that had proliferated many times and had expended 60-70% of their lifespan prior to culture. In contrast, TSC which can be passaged 15-20 times are a subset of TSC that proliferated fewer times in vivo and had expended only 30-40% of their lifespan. Thus, differences in passage-potential reflect the TSC’s contribution to epithelial homeostasis and repair prior to isolation.

Our finding that TSC frequency plateaus after passage 5 suggested the presence of an additional TSC conservation mechanism. Although some studies reported that TERT was active (Yim, Slebos et al. 2007, Suprynowicz, Upadhyay et al. 2012), our RNAseq studies did not identify a population of TERT expressing TSC (Fig 6D, SFigs 4 and 5). However, we did detect expression of shelterin genes in TSC and our analysis of TSC from DC patients and carriers suggested that shelterin complexes may maintain the relatively short telomeres found in long-lived TSC (Fig 7). Finally, our analysis of secretory specification in TSC (Fig 6F-G, SFig 6) complements our analysis of terminal differentiation in mouse TSC clones (Fig 3) and suggests that asymmetric cell division may maintain the TSC population in long-lived clones.

The “TSC exhaustion” concept suggests that compromised TSC function leads to airway epithelial remodeling in chronic lung disease. This process may be due to a change in TSC differentiation potential in healthy smokers and smokers with COPD (Ghosh, Miller et al. 2018) but could also include activation of the terminal differentiation pathway (Figs 3, 6). A role for telomere biology in TSC exhaustion and chronic lung disease is suggested by identification of TERT risk alleles in idiopathic pulmonary fibrosis (Armanios and Blackburn 2012) and our demonstration of telomere shortening in human TSC (Fig 6) and in DC families (Fig 7). Collectively, these studies identify biological aging as an important aspect of TSC phenotype and function and one that could impact the development of chronic lung disease.

## ACKNOWLEDGEMENTS

This project was funded by Research Grants the Cystic Fibrosis Foundation (SDR), the Nationwide Children’s Hospital Cellular Therapy and Cancer Immunotherapy Program (SDR, DH), and the Cystic Fibrosis Foundation Cure Columbus Research Development Program (SDR, DH). RO1, HL129938 (MG), and FAMRI grant 113259_YCSA (MG).

## AUTHOR PARTICIPATION

MG: Conception and design, acquisition of data, interpretation of data, manuscript preparation and review

CLH: Acquisition of laboratory data, manuscript review AA: Acquisition of laboratory data, manuscript review SWL: Acquisition of laboratory data, manuscript review

JEM: Conception and design, acquisition of data, interpretation of data, manuscript review DH: Acquisition of clinical samples, manuscript review

SW: Conception and design, acquisition of data, interpretation of data, manuscript review SDR: Conception and design, acquisition of laboratory data, interpretation of data, manuscript preparation and review.

## FIGURE LEGENDS

**SFig 1. Related to Fig 2.**
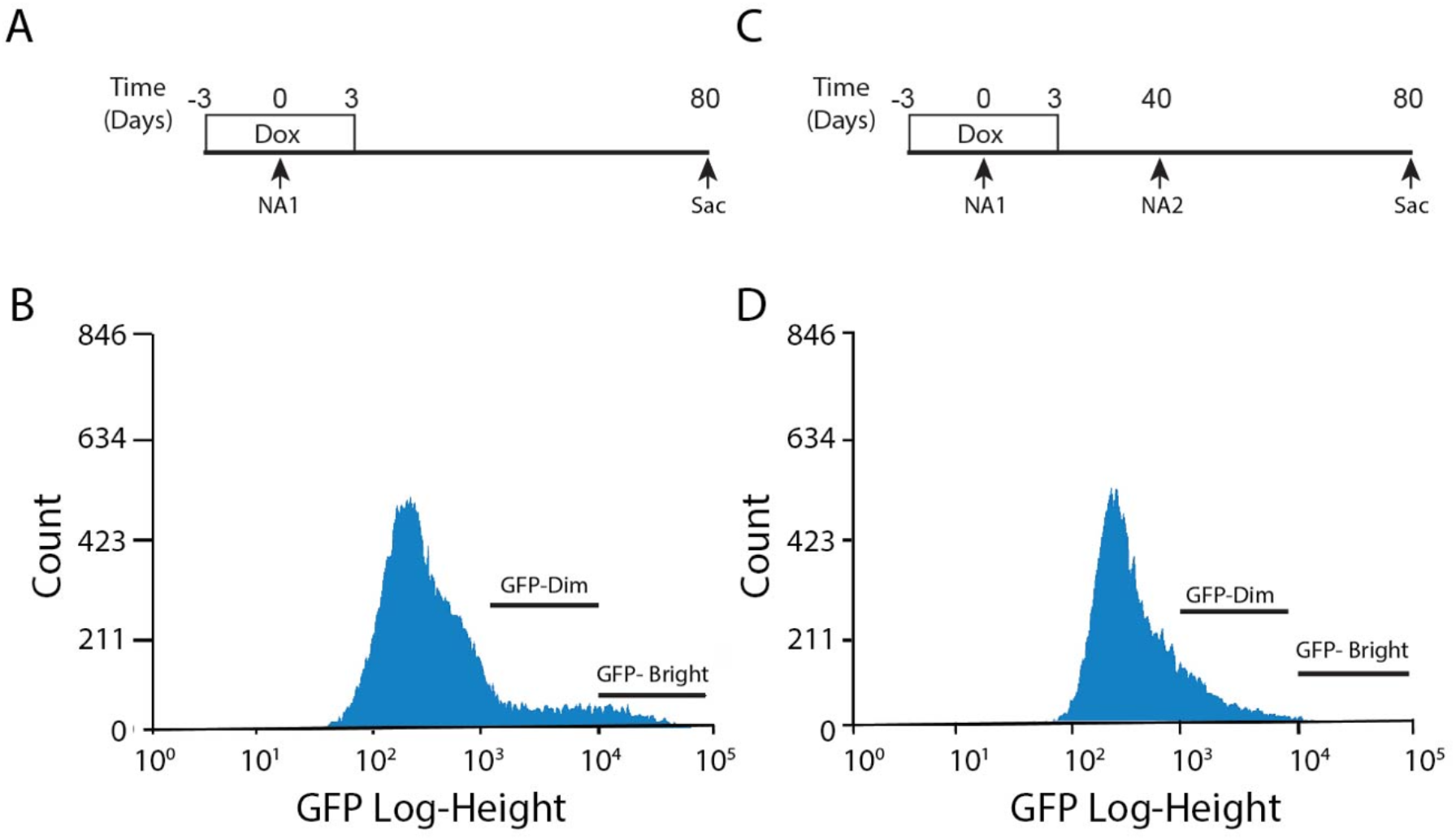
A second NA injury accelerates TSC proliferation. All studies used K5/H2B:GFP mice and mono-transgenic controls. **A**. Mice were treated with Dox chow from day −3 through day +3 and then switched to standard chow. NA treatment occurred on day 0. Animals were euthanized (sac) on day 80. **B**. Flow cytometry was used to evaluate green fluorescent protein (GFP) fluorescence intensity 80 days after NA1. **C**. Mice were treated Dox and NA as indicated above, retreated with NA on day 40 (NA2), and euthanized on day 80. **D**. Flow cytometry was used to evaluate GFP fluorescence intensity 40 days after NA2. Representative of 3 biological replicates.

**SFig 2. Related to Fig 3.**
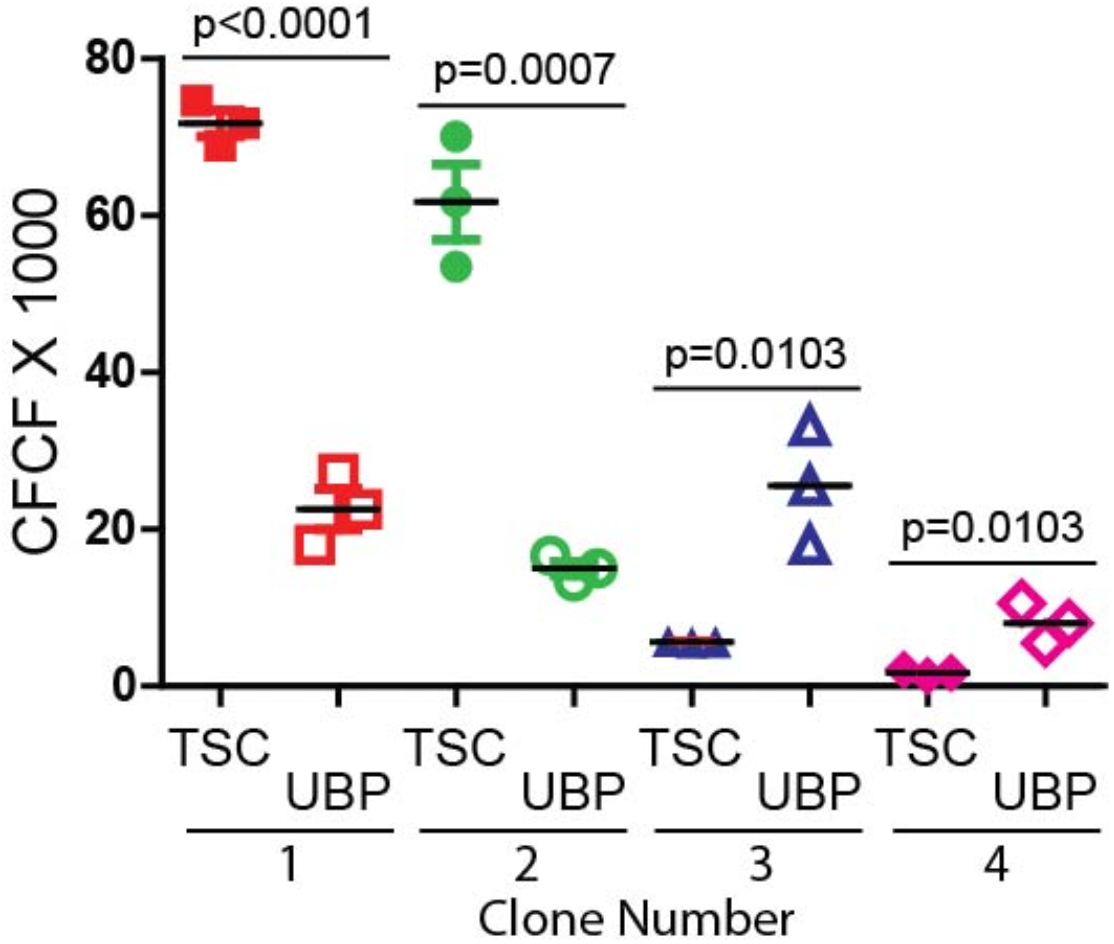
Analysis of TSC and UPB frequency in TSC clones. TSC were recovered from mice that were injured with NA and recovered 40 days. The clone forming cell frequency (CFCF) assay was used to determine the fate of 4 clonal isolates (clone numbers 1-4) at passage 3. Solid symbols-TSC clones; open symbols-UPB clones; red symbols-clone 1, green symbols-clone 2; blue symbols-clone 3; pink symbols-clone 4. Mean ± SD (n=3).

**SFig 3. Related to Fig 4.**
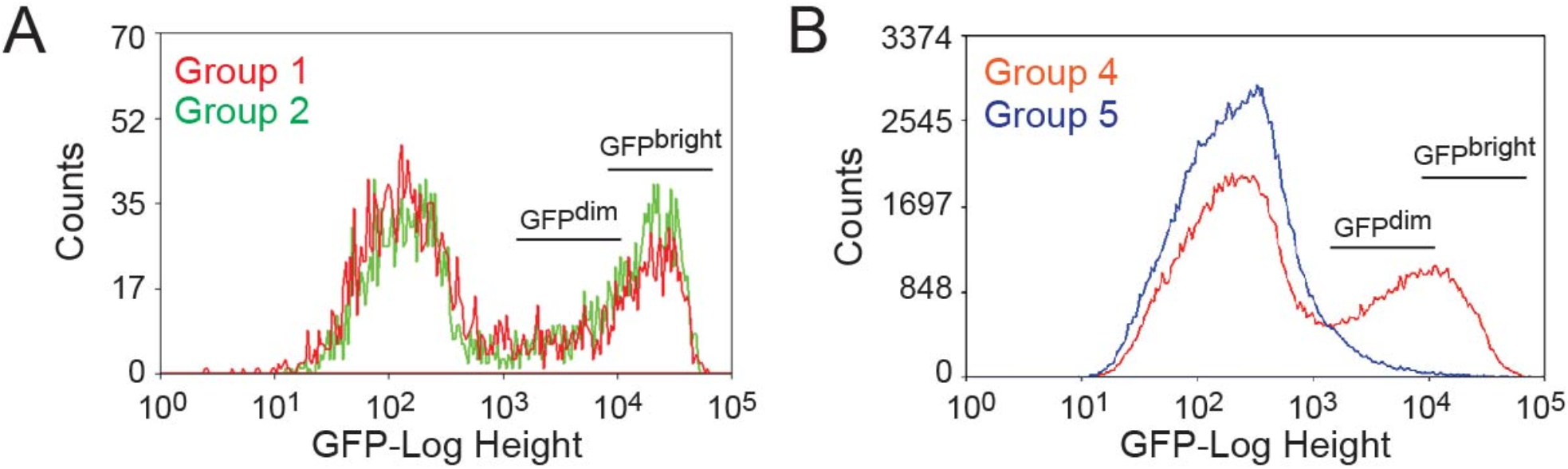
See Fig 4 for experimental design. **A**. Flow cytometric analysis of green fluorescent protein (GFP) fluorescence intensity 6 days after the first or second NA injury. Representative of 3 studies. **B**. Flow cytometric analysis of green fluorescent protein (GFP) fluorescence intensity 40 days after the first or second NA injury.

**SFig 4. Related to Fig 6.**
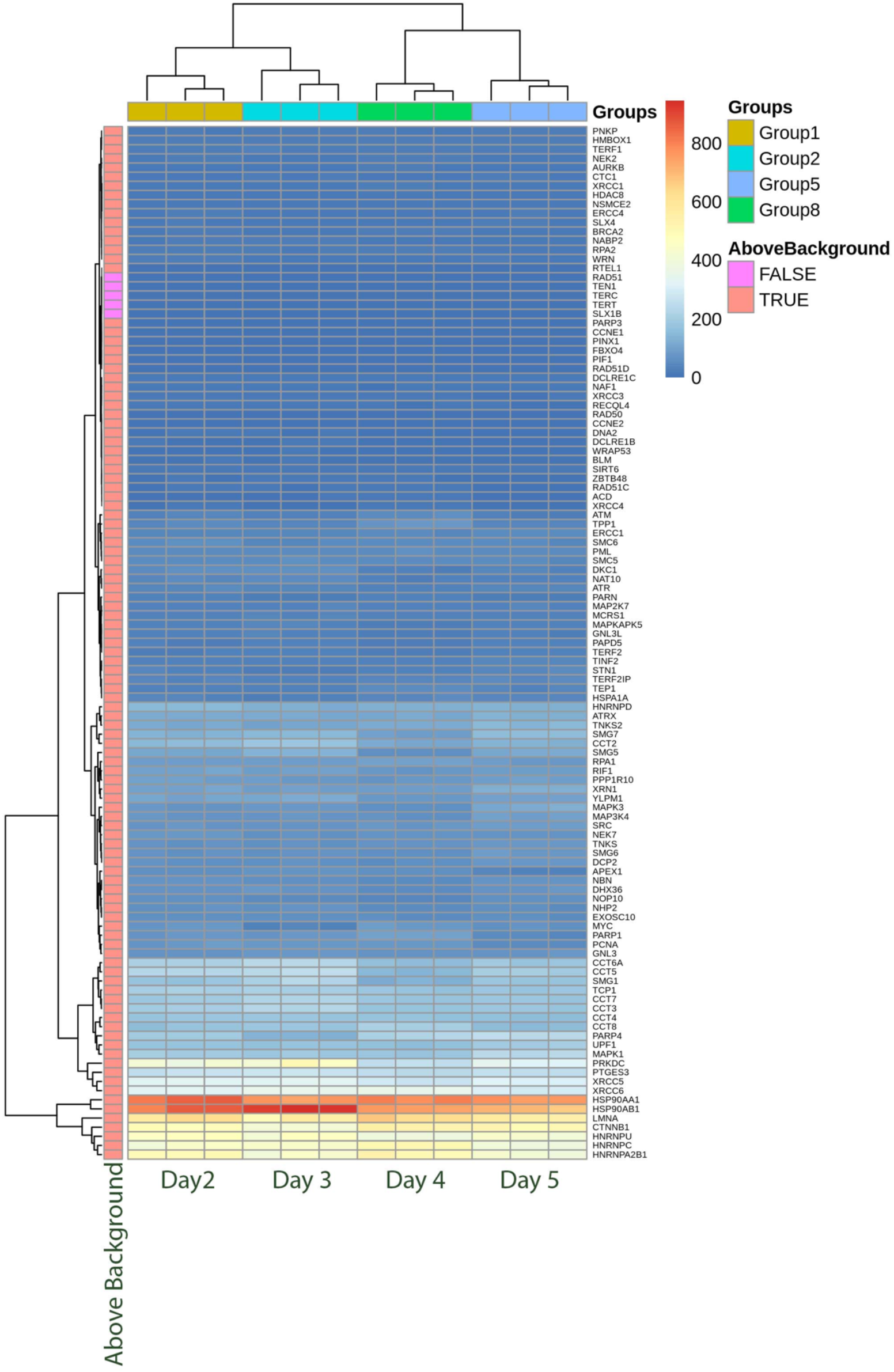
Large scale presentation of a heatmap representing telomere biology gene expression in TSC at passage 2. RNAseq analysis on post-plating days 2, 3, 4, and 5 (n=3). Gene expression, in Fragments Per Million Reads (FPM), for telomere biology genes is shown. Background (BKG): FPM > 0.5 = above background (salmon), FPM<0.5 below background (plum including TERT and TERC).

**SFig 5. Related to Fig 6.**
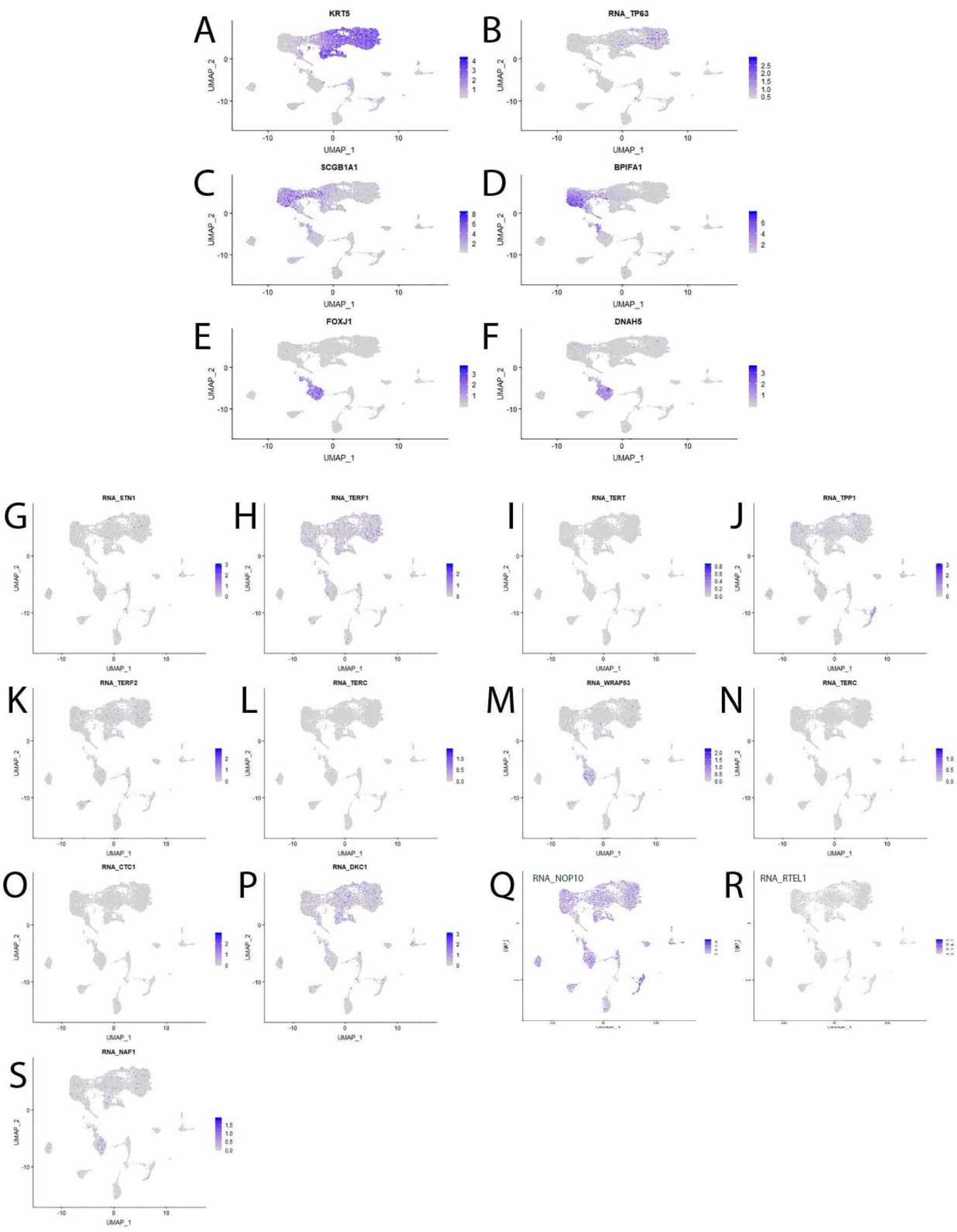
Telomere biology gene expression freshly isolated human lung cells. Single cell RNAseq was used to evaluate gene expression. **A-F**. Feature maps illustrating the distribution of Keratin 5+ (A), and TRP63+ (B) basal cells; SCGB1A1+ (C) and BPIFA1+ (D) secretory cells; and FoxJ1+ (E) and DNAH5+ (F) ciliated cells. **G-O**. Feature maps illustrating the distribution of cells expressing various telomere biology genes. Gene names are indicated in each panel. TERT and TERC gene expression was not above background.

**SFig 6. Related to Fig 6.**
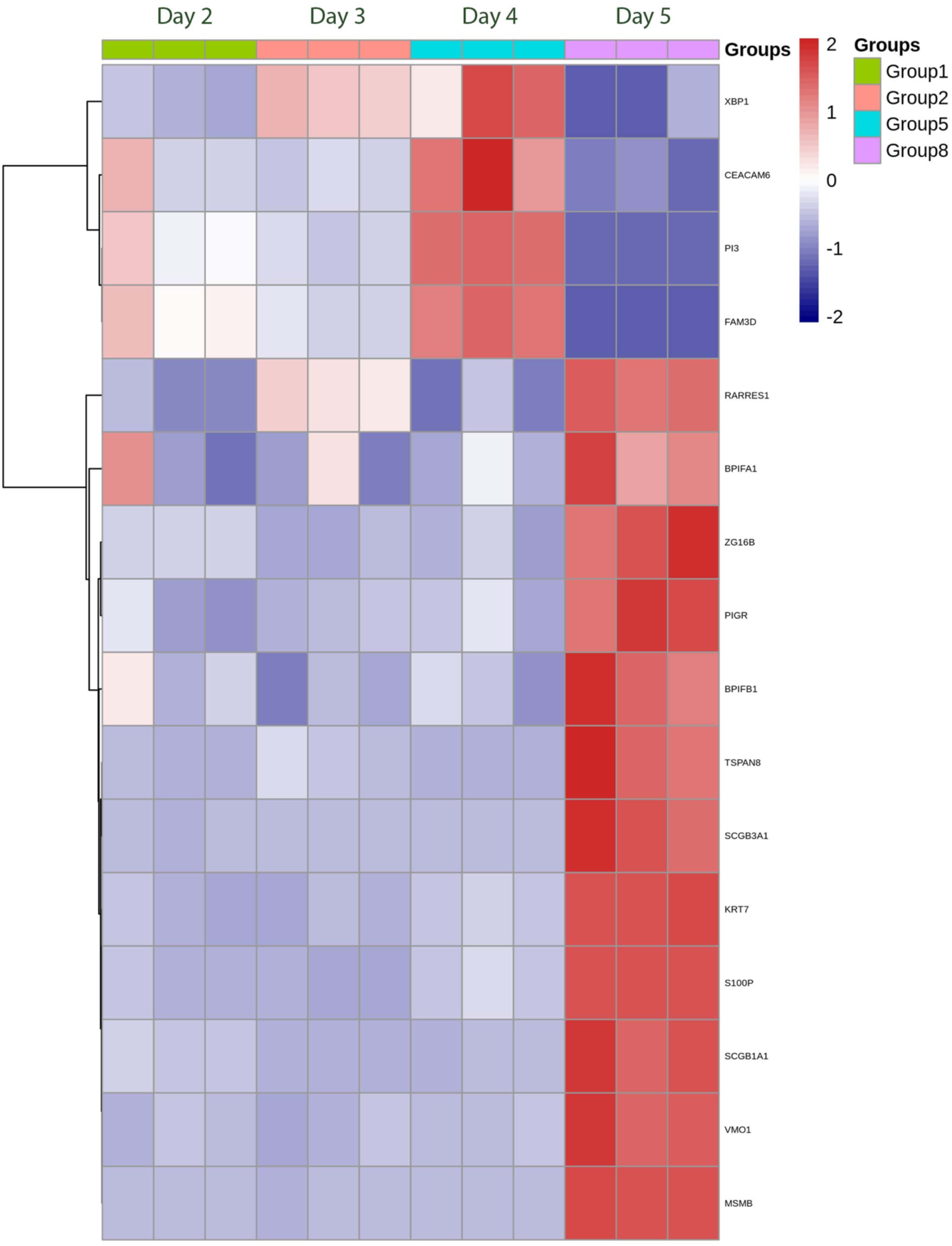
A heatmap representing secretory gene expression in TSC at passage 2. RNAseq analysis on post-plating days 2, 3, 4, and 5 (n=3). Gene expression is presented in rows as the scaled z-score. For an example, a value of 2 means gene expression is 2 standard deviations above the mean for the given row. This format provides a clearer representation of changes in the expression of cell type-specific signature genes than the fragments per million reads approach.

**STable 1.** Donor demographics for dyskeratosis congenita probands and parents.

| Donor | Age | Sex | Mutation | Pulmonary | Bone Marrow |  | Liver | GI Tract | Bone | Mouth |
| --- | --- | --- | --- | --- | --- | --- | --- | --- | --- | --- |
|  |  |  |  |  | Failure | Transplant |  |  |  |  |
| 1 | 15 | M | RETL1 | AVM | 1 | 1 | 0 | 0 | HAN | LP |
| 2 | 12 | M | TPP1 | AVM, PF | 1 | 1 | 0 | ES | OPO | LP |
| 3 | 55 | F | TERT | PF | 0 | 0 | FL | GER, GP, IBS | HAN, OPO, SF | GD |
| 4 | 26 | M | WRAP53 | 0 | 1 | 1 | 0 | ES | KAN | LP |
| 5 | 19 | M | RTEL1 | AVM | 1 | 1 | 0 | ES | HAN | LP |
| 6 | 13 | M | TERT | Abn CXR | 0 | 0 | 0 | GER | 0 | 0 |
| 7 | 58 | F | RTEL1 | Low DLCO | 0 | 0 | 0 | 0 | OPA, FR | 0 |
| 8 | 41 | F | TINF2 | 0 | 0 | 0 | 0 | 0 | OA | LP |
| 9 | 40 | M | U | 0 | 0 | 0 | HS, NRH | 0 | 0 | 0 |
| 10 | 29 | M | CTC1 | 0 | 0 | 0 | LC | 0 | FR | GD |
| 11 | 5 | M | TINF2 | 0 | 1 | 1 | 0 | GER, ES | 0 | LP |
| 12 | 18 | M | U | Asthma | 0 | 0 | 0 | GER | OPO | FC |

0-absent; 1-present; AVM-arteriovenous malformations; Abn CXR-abnormal chest x-ray; DLCO-diffusing capacity for carbon monoxide; ES-esophageal structure; F-female; FC-frequent cavities; FL- fatty liver; FR-fractures; GD-gum disease; GER-gastroesophageal reflux; GI-gastrointestinal; GP-gastroparesis; HAN-hip avascular necrosis; HS-hepatopulmonary syndrome; IBS-inflammatory bowel syndrome; KAN-knee avascular necrosis; LC-liver cirrhosis; LP-leukoplakia; M-male; NRH-nodular regenerative hyperplasia; OA-osteoarthritis; OPA-osteopenia; OPO-osteoporosis; PF-pulmonary fibrosis; SF-stress fractures; U-unknown.

## Methods

### Resource Availability

#### Lead Contact

Further information and requests for resources and reagents should be directed to and will be filed by the Lead contact, Susan D. Reynolds

#### Materials Availability

This study did not generate new unique reagents.

#### Data and Code Availability

Gene expression data will be made available at GEO.

### Experimental Model and Subject Details

#### Mouse models

All procedures involving animal use were approved by the Institutional Animal Care and Use Committee at National Jewish Health or Nationwide Children’s Hospital. Mice were maintained in an AAALAC-approved facility and screened quarterly for pathogens. All mice were research naïve. Mice were group housed, fed ad libitum, and maintained on a 12 hour light/dark cycle. Littermates of the same sex were randomly assigned to experimental groups.

#### Human studies

The Institutional Review Board at Nationwide Children’s Hospital approved the human studies. Written informed consent and assent was obtained from every participant.

Normal donor demographics were previously reported (Hayes). Bronchial tissue samples were obtained at the time of lung transplantation. Dyskeratosis congenita donor age, sex, and genotype are summarized in Supplemental Table 1.

### Method Details

#### Naphthalene (NA) exposure

NA was prepared and administered by intraperitoneal injection as previously reported (Cole, Smith et al. 2010). NA dose was selected to cause >95% depletion of the Club cell secretory protein (CCSP) positive cell population by recovery day 3.

#### Chromatin labeling

The Keratin 5-reverse tetracycline trans-activator/tetracycline responsive element-Histone 2B:green fluorescent protein (K5-rtTA/TRE-H2B:GFP) chromatin-labeling model facilitates purification of basal cells that proliferate frequently or infrequently after injury (Tumbar, Guasch et al. 2004). Adult male and female K5-rtTA (Diamond, Owolabi et al. 2000)/TRE-histone 2B mice were used for the label-retention studies. Adult (6-12 weeks old) male and female mice were used. Genetic background was a mixture of C57Bl/6 and FVB/n. Body weight was 25-40 grams at the time of treatment. Mice were fed doxycycline (Dox) chow and treated with 250 mg/kg NA according to the schedule detailed in Results.

Adult female (6-8 weeks old) inbred FVB/n mice were treated with 300 mg/kg NA. Body weight was 22-28 grams at the time of treatment. Groups of mice were treated with 50 mg/kg Bromodeoxyuridine BrdU and/or 42 mg/kg 5-ethynyl-2’-deoxyuridine (EdU) according to the schedule detailed in Results.

#### Immunofluorescence studies

Tracheal tissue sections (5 μm) were generated from paraffin-embedded tissues and processed as described previously (Smith, Hicks et al.). All antibodies (KRT5, CCSP, BrdU, and ACT), and staining methods were previously described (Cole, Smith et al. 2010). EdU was detected using the ClickIt method and followed the manufacturer’s instructions. Images were acquired using an upright Zeiss Imager Z1 fluorescent microscope and AxioVision software (Carl Zeiss) or an inverted Zeiss 200M confocal microscope using Intelligent Imaging Innovations Inc. software. Cell type frequency was quantified as previously indicated (Smith, Hicks et al.) and values were expressed as a percent of total epithelial cells. N=6 tracheas were quantified/group.

#### Mouse TSC quantification

Tracheal epithelial cells were recovered by dispase/collagenase/ trypsin (DCT) digestion and clone forming cell frequency (CFCF) was determined by limiting dilution (Ghosh, Helm et al. 2011). Mouse cells were plated on irradiated NIH3T3 (ATCC #CRL-1658) feeder layers cultured in mouse tracheal epithelial medium-plus (MTEC+, (You, Richer et al. 2002)). On culture day 10, the cells were fixed and stained with Geimsa. TSC clones were identified as previously reported (Ghosh, Helm et al. 2011). Time points are noted in Results.

#### Mouse flow cytometry

Tracheal epithelial cells were recovered by digestion with DCT. A Moflo high-speed cell sorter (Dako Cytomation) was used. First, cells that were CD45-/CD31-/TER119-/DAPI-were excluded. Next, CD49f^bright^/Sca1+ cells were identified and GFP fluorescence intensity in each cell was quantified. GFP^bright^ cells were defined as those having a log GFP fluorescence >10^4^, GFP^dim^ cells were defined as those having a log GFP fluorescence of 10^3^-10^4^. GFP^bright^ and GFP^dim^ cells were sorted into MTEC+ medium.

#### Mouse serial passage analysis

Passage (P) 0 TSC clones were derived from NA treated mice that had been recovered 80 days. One thousand tracheal epithelial cells were plated in each well of a 6-well plate that contained an irradiated NIH3T3 fibroblast feeder cell layer. The cells were cultured in MTEC+ medium for 10-days. TSC clones were identified by their characteristic rim morphology and isolated using cloning cylinders as previously reported (Ghosh, Helm et al. 2011). This process was repeated at P3. At P5, the number of cells/clone and the TSC- and UPB-CFCF were used to calculate the number of TSC and UPB cells.

#### Irradiated NIH3T3 feeder layers

NIH3T3 fibroblasts (ATCC #CRL-1658) were recovered a mus musculus (mouse) embryo. The strain was NIH/Swiss. Sex is not known. Following receipt, the cells were cultured and crypreserved in multiple aliquots. Cells were not further characterized. NIH3T3 cells were cultured in DMEM, 10% fetal bovine serum, L-glutamine, and penicillin/streptomycin at 37 °C in 5% CO2 for up to 10 passages. NIH3T3 cells recovered with trypsin/EDTA and either subcultured or irradiated with 2500 Grey using a X-irradiator. Irradiated cells were plated at 3.2 × 10^5^ cells/cm^2^ in as indicated above and allowed to adhere overnight.

#### Human TSC recovery and culture

Human bronchial TSC were recovered as previously described (Reynolds, Rios et al. 2016, Hayes, Kopp et al. 2018). The modified conditional reprogramming culture (mCRC) method (Reynolds, Rios et al. 2016) was adapted from Suprynowicz et al (Suprynowicz, Upadhyay et al. 2012). The major change from this protocol was the use of irradiated NIH3T3 fibroblast feeder layers (see below).

#### Human TSC enumeration

TSC were enumerated using the limiting dilution assay as previously reported (Ghosh, Ahmad et al.). The CFCF assay (Ghosh, Helm et al. 2011) was used to determine TSC number.

#### Human TSC cloning

TSC were cloned at P3 using the limiting dilution method (Hayes). Passage 1 basal cells were seeded at 5 cells/well in 96-well cell culture and cultured according to the mCRC method for 5-7 days. Wells containing a single colony were identified and verified after removal of the fibroblast feeder layer. The basal cells were then recovered by a second trypsinization and cultured in one well of a 12-well plate.

#### TSC differentiation

TSC were plated onto collagen-coated 0.33 cm^2^ transwell membranes at 2×10^4^ cells per membrane as previously described (Reynolds, Rios et al. 2016). At confluence, the medium was changed to Half and Half differentiation medium (Malleske et al, in press) which was composed of equal volumes of Wu differentiation medium (Wu 2000) and Pneumacult Base Medium containing the 10X supplement, heparin, and hydrocortisone (Stemcell Technologies, Vancouver, BC, Canada). On differentiation day 21, the cultures were fixed and stained. Nuclei were detected using 4′ 6-diamidino-2-phenylindole (DAPI), mucus cells were detected with rabbit-anti-MUC5b (1/100, (Seibold, Smith et al. 2013)), and ciliated cells were detected with mouse-anti-acetylated tubulin (1/8000, ACT, (Seibold, Smith et al. 2013)).

Differentiation was quantified using a serological method (Reynolds, Rios et al. 2016) that meets the American Thoracic Society standard for assessment of histological data sets (Hsia, Hyde et al. 2010).

#### Telomere length determination

Genomic DNA was purified using the DNeasy Blood & Tissue Kit (69506, Qiagen); 40ng of gDNA was used for telomere and 36B4 single copy gene control assays. Targets were amplified using previously reported standards and primers (O’Callaghan, Dhillon et al. 2008). Previously published cycling parameters for telomeres (Hsieh, Saberi et al. 2016) and 36B4 (O’Callaghan, Dhillon et al. 2008) were used. Magnesium concentration in the PowerSYBR Green Master Mix (Applied Biosystems 4367659) was lowered through the addition of 0.5M EDTA (0.0385µl per 20µl reaction for a final concentration of 0.9625mM) for the telomere assay (Hsieh, Saberi et al. 2016). Jurkat (ATCC) and U1301 (human T-cell leukemia cell line (01051619, Sigma) were used as short and long telomere controls, respectively. A standard curve was generated for each experiment and was used to insure a linear relationship between DNA input and telomere length. Three technical replicates were generated for each standard curve point and biological sample. Relative telomere length was calculated according to the Telomere/Single copy gene control (T/S) ratio (O’Callaghan, Dhillon et al. 2008). The XX assay was used to independently determine telomere length in a subset of samples.

#### RNA-Seq data analysis

On average, 32 million paired-end 151 bp RNA-Seq reads were generated for each sample (the range was 25 to 37 million). Each sample was aligned to the GRCh38.p9 assembly of the Human reference from NCBI (http://www.ncbi.nlm.nih.gov/assembly/GCF_000001405.35/) using version 2.6.0c of the RNA-Seq aligner STAR (http://bioinformatics.oxfordjournals.org/content/29/1/15). Transcript features were identified from the GFF file provided with the assembly from Gencode (v28) and raw coverage counts were calculated using featureCounts (http://bioinf.wehi.edu.au/featureCounts/). The raw RNA-Seq gene expression data was normalized and post-alignment statistical analyses were performed using DESeq2 (http://genomebiology.com/2014/15/12/550) and custom analysis scripts written in R. Comparisons of gene expression and associated statistical analysis were made between different conditions of interest using the normalized read counts. All fold change values are expressed as test condition / control condition, where values less than one are denoted as the negative of its inverse (note that there will be no fold change values between –1 and 1, and that the fold changes of “1” and “-1” represent the same value).

Transcripts were considered significantly differentially expressed using a 10% false discovery rate (DESeq2 adjusted p value <= 0.1). Single cell RNAseq methods and analytical techniques are presented in detail in (Carraro, Mulay et al. 2020).

### Quantification and Statistical Analyses

All statistical analyses were performed using Graph Pad Prism. Outliers were identified by the ROUT test. Data normality was evaluated by the Shapiro-Wilk test, the Anderson-Darling test, and/or the D’Agostino & Pearson test. For normally distributed data sets, differences were evaluated using Student’s t-test and data are presented as the mean ± standard deviation or the standard error of the mean. For non-normally distributed data sets, differences were evaluated by the Mann-Whitney test and data are presented as the median and the interquartile range.

Trends were analyzed by regression analysis. Data sets containing multiple variables were analyzed by analysis of variance (ANOVA) and a post hoc Tukey test (normally distributed data sets) or Kruskal Willis test (non-normally distributed data sets). Data are presented as indicated above. The specific statistical method, sample size, definition of center, and dispersion precision measures are indicated in the Figure Legends. P-values < 0.05 were considered to be significant.

### Additional Resources

Not applicable

## KEY RESOURCES TABLE

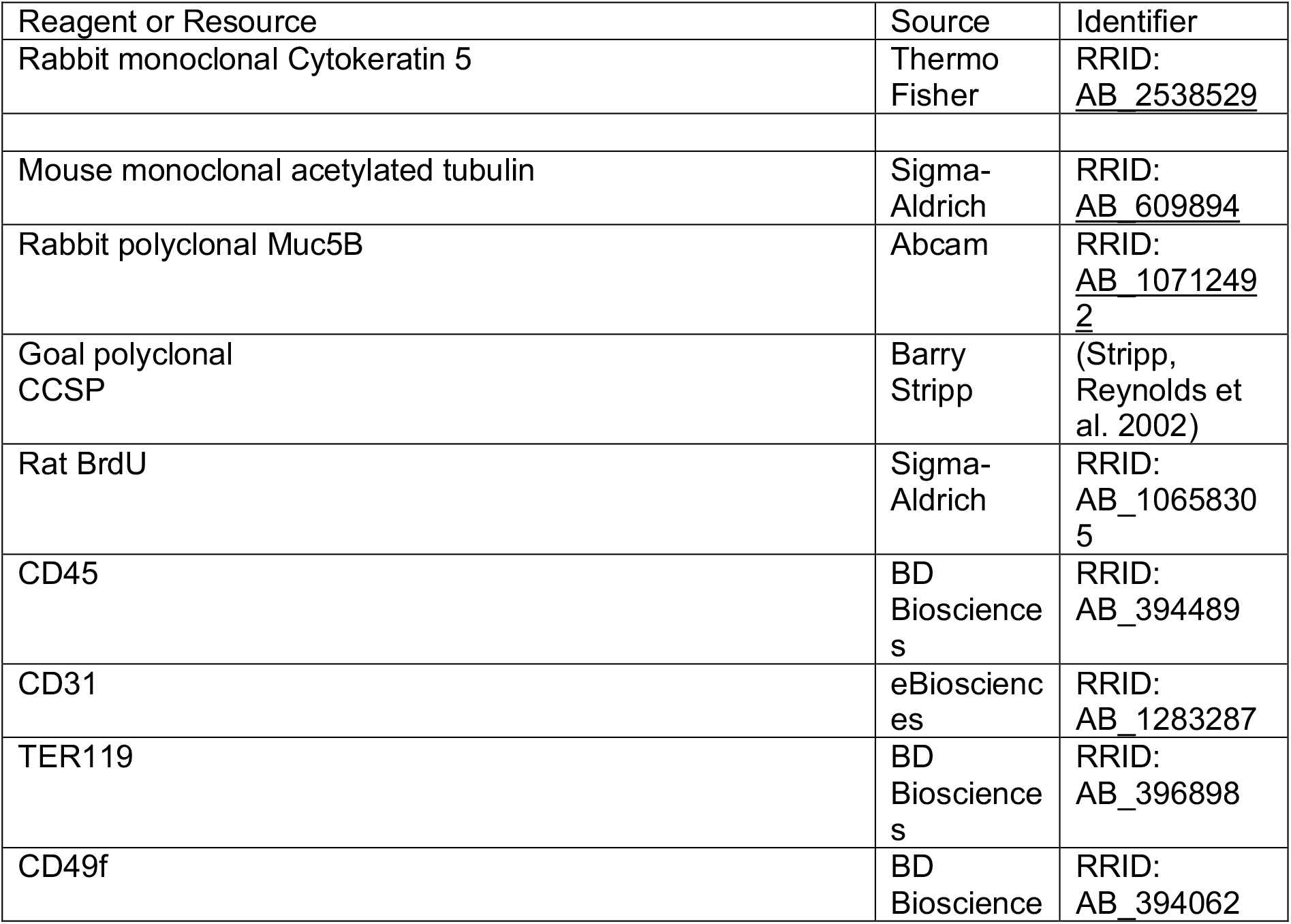
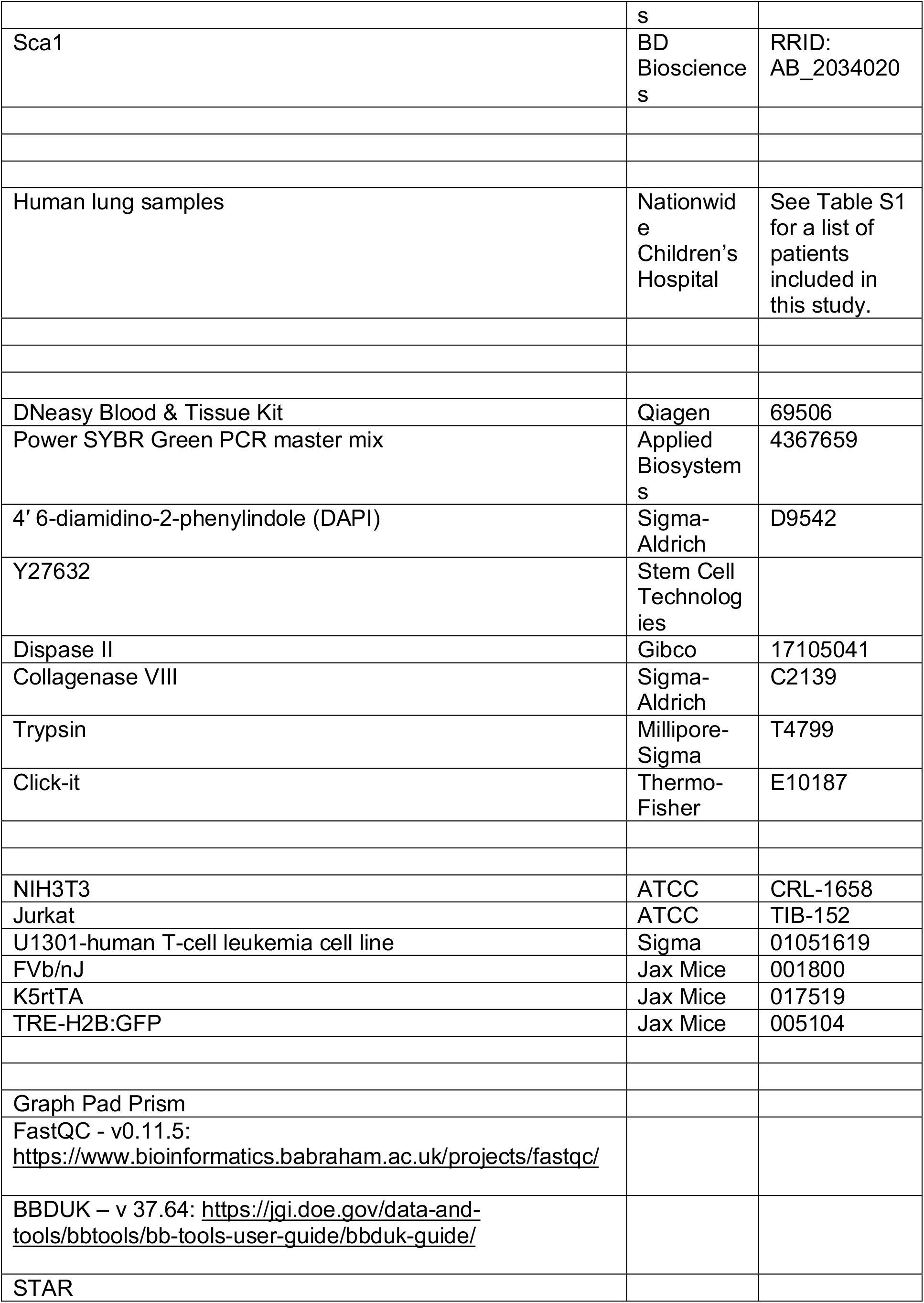
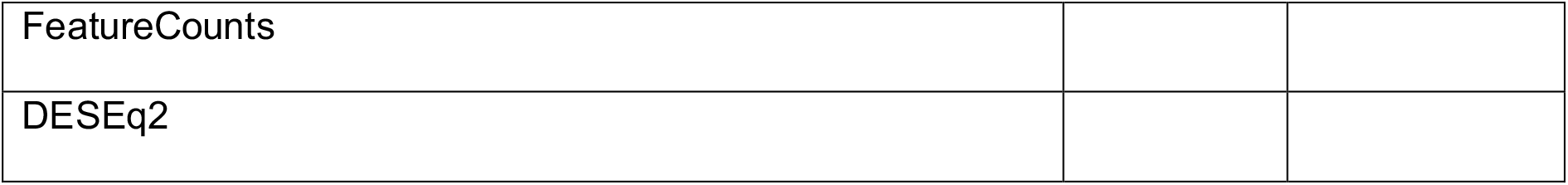

## Supplemental data

